# Unlocking the potential of *Uvaria chamae*: A cream formulation for combatting *Staphylococcus aureus* skin infections

**DOI:** 10.1101/2024.06.28.601194

**Authors:** Eskyl Togbé, Hornel Koudokpon, Aimé Césaire Ayéna, Phénix Assogba, Edna Hounsa, Amy Watara, Eric Agbodjento, Fréjus Ohouko, Jean Robert Klotoé, Lamine Baba-Moussa, Victorien Dougnon

## Abstract

*Staphylococcus aureus* (*S. aureus*) skin infections remain a prevalent public health concern worldwide, posing ongoing challenges in treatment. *Uvaria chamae* roots has demonstrated significant effectiveness against *S. aureus*. This study aimed to propose a dermal formulation based on *Uvaria chamae* for treating bacterial skin infections. The antibacterial activity was first confirmed. Tests including the time-kill test, the erythromycin outer membrane permeability test, and the ATPase/H+ proton pump inhibition test were conducted to explore the antibacterial mode of action. A cream was prepared using the phase inversion technique and subjected to quality control and senso-rheological testing. The cream was tested for antibacterial activity and skin toxicity (OECD guideline 404). The *in vivo* antibacterial effect and wound-healing potential of the *Uvaria chamae* cream were then assessed on Wistar rat. The ethanolic extract of *Uvaria chamae* root exhibited bactericidal effects on the tested *S. aureus* strains. In combination with erythromycin, the extract enhanced the permeability of *S. aureus*’s outer membrane while inhibiting ATPase/H+ proton pumps. The pharmaceutical quality of the produced cream revealed acceptable macroscopic conditions and satisfactory homogeneity. Antibacterial activity evaluation showed significant activity at low concentrations. The cream’s healing properties indicated a significant decrease in wound diameter (p<0.05) over time, culminating in complete healing on day 13 for the treated group, with a retraction percentage of 99.99%. These findings support the safe use of *Uvaria chamae* extracts in managing *S. aureus* skin infections and pave the way for pre-clinical testing and potential market introduction of this cream.

## Introduction

Skin infections remain one of the most widespread public health problems globally, continually presenting treatment challenges [1]. Bacterial infections, a common type of dermatological condition, result from the proliferation and harmful activity of bacteria on the skin [2,3]. Notably, *S. aureus* and *Streptococcus pyogenes* are frequent culprits behind these infections [4,5].

The treatment of bacterial skin infections depends on the infection type, severity, and the bacteria involved. Often, topical or oral antibiotics are administered to eliminate the responsible bacteria [6]. However, the inappropriate and uncontrolled use of antibiotics has led to a concerning increase in antimicrobial resistance, resulting in therapeutic failures, more severe infections, and higher healthcare costs, thus posing a significant public health issue [7]. The rising prevalence of antibiotic resistance among skin pathogens underscores the urgent need to discover new antimicrobial compounds [1]. Plants are emerging as one of the most promising sources for new antibacterial agents [8,9]. Consequently, recent years have seen extensive research into the antimicrobial activity of African flora, particularly the flora of Benin. Among the most frequently cited plants for their antimicrobial properties is *Uvaria chamae* [10]. *Uvaria chamae,* a plant with notable medicinal properties, is becoming a potential source for new antimicrobial solutions [11]. Common in tropical Africa, its edible fruits and roots are renowned worldwide for their medicinal, cosmetic, and nutritional properties [12]. Several studies have identified various bioactive compounds in this plant that exhibit antimicrobial properties against numerous pathogenic microorganisms, including antibiotic-resistant bacteria, thus broadening its therapeutic applications [13]. Research has shown that ethanolic extracts of *Uvaria chamae* roots contain chalcones and dihydrochalcones, bioactive compounds that impart antimicrobial activity against a range of multi-resistant gram-positive bacteria at lower concentrations than conventional antibiotics [11,14]. Consequently, *Uvaria chamae*, with its well-established medicinal properties, emerges as a promising candidate for treating skin infections. Nonetheless, a detailed understanding of its mode of action and antimicrobial potential, particularly concerning skin infections, warrants further investigation [1].

In phytotherapy for skin infections, various preparation methods, including infusions and poultices, are being explored [15]. While poultices can be beneficial, they also pose risks, such as phytophotodermatoses [16]. Therefore, developing a *Uvaria chamae*-based cream is a commendable initiative, presenting low risks and offering a practical, easy-to-apply formulation that is compatible with the skin. This context forms the basis of the present study, which aimed to propose a dermal formulation based on *Uvaria chamae* for treating *S. aureus* skin infections.

## Materials and Methods

### Ethical consideration

The research protocol received approval from the Ethics Committee of the Research Unit in Applied Microbiology and Pharmacology of Natural Substances at the University of Abomey-Calavi in Benin (Approval No. 0043/2022/CE/URMAPha/UAC). All procedures were carried out following the guidelines of the National Institute of Health (NIH) for the care and use of laboratory animals.

### Collection, identification, and processing of plant material

Fresh *Uvaria chamae* roots were sourced from herbalists in the Abomey-Calavi commune and authenticated at the National Herbarium of the University of Abomey-Calavi (NH-UAC) under the number AA6687/HNB. Once authenticated, the roots were dried at room temperature in the Research Unit of Applied Microbiology and Pharmacology of Natural Substances (URMAPHA) at the University of Abomey-Calavi (UAC). The dried roots were then ground into a powder using a dedicated mill.

### Animal material

Female Wistar albino rats weighing between 180 and 200 g were used. These rats came from the animal house of the Applied Microbiology and Pharmacology of natural substances research unit. They were randomly assigned to standard cages with ad libitum access to food and water. Throughout the study, the rats were housed in an environment maintained at a constant temperature of 22°C, with a regular 12-hour light/dark cycle.

### Microorganisms

Five strains of Gram-positive cocci and five strains of Gram-negative bacilli, including both reference and clinical strains, were provided by the URMAPHA at the UAC. These strains included *Escherichia coli* ATCC 25922, *Escherichia coli* Extended-Spectrum Beta-Lactamases (ESBL), *Klebsiella pneumoniae* ESBL, *Salmonella* spp, *Pseudomonas aeruginosa*, *S. aureus* ATCC 25923*, S. aureus* ATCC 6835, cefoxitin-resistant *S. aureus*, MecA-resistant *S. aureus*, and vancomycin-resistant *Enterococcus faecalis*.

### Extract preparation

The study utilized ethanolic extracts based on the study of Koudokpon et al. [11, 17] on Gram-positive cocci. The extract was prepared following the protocol by Koudokpon et al. [11,17]. Briefly, 200 g of *Uvaria chamae* root powder was dissolved in 2000 ml of 96° ethanol and continuously stirred for 72 hours. The mixture was then filtered through No. 2 and 4 filter paper. The filtrate underwent dry evaporation using a Rotavapor at 600 rpm at 45°C, and the remaining mixture was oven-dried at 45°C for 24 hours to yield a dry extract. This extract was stored at 4°C and reconstituted for various tests.

### Direct *in vitro* antibacterial activity of *Uvaria chamae* hydroethanolic extract

A 0.5 McFarland bacterial suspension from an 18-hour culture of each strain was prepared according to CASFM guidelines [18]. Each suspension was swabbed onto Mueller-Hinton agar, and two 6 mm-diameter wells were drilled using the sterile tip of a Pasteur pipette. One well received 50 μl of the sterile extract solution, while the other received a 10% DMSO solution as a negative control. Positive controls included conventional antibiotic discs of vancomycin (5 µg) and ciprofloxacin (5 µg) for Gram-positive cocci, and imipenem (10 µg) and fosfomycin (200 µg) for Gram-negative bacilli. Petri dishes were left at room temperature for one hour for pre-diffusion, then incubated at 37°C for 18 hours. Inhibition diameters were measured and compared to standards [19]

### Determination of Minimum Inhibitory Concentration (MIC) and Minimum Bactericidal Concentration (MBC)

MIC determination followed the microwell method described by Dougnon et al. [20]. Each well received 100 µl of the initial extract solution (5 mg/ml) and 100 µl of Mueller-Hinton broth, followed by serial dilution. After adding 100 µl of bacterial suspensions, microplates were incubated at 37°C for 18 hours. Tetrazolium solution was added, and after 20 minutes of dark incubation, red coloration indicated viable bacteria. MIC was the lowest concentration with no red wells. Non-red wells were plated on Mueller-Hinton agar and incubated at 37°C for 24 hours to determine BMC. The BMC/MIC ratio indicated whether the extract was bacteriostatic (≥4) or bactericidal (<4).

### Kinetic Time-Kill Test

The kinetic study of the ethanolic extract was conducted using the method described by Djague et al. [21] with modifications. Bacterial suspensions at various MIC multiples (1x, 2x, 4x) were incubated at 37°C, and optical density was measured at 600 nm at intervals (0, 2, 4, 6, 8, 24 hours). Ciprofloxacin served as the positive control, and MHB with bacterial strain as the growth control. The test was repeated three times, and growth curves were plotted.

### Erythromycin outer membrane permeability test

The effect on outer membrane permeability was assessed using the method by Ekom Tamokou and Kuete et al. [22] with modifications. In a 96-well microplate, 50 µl of Mueller-Hinton broth, 50 µl of erythromycin solution, and various concentrations of the ethanolic extract were combined with 50 µl of bacterial suspension. After 24 hours of incubation at 37°C, bacterial growth was measured at 450 nm.

### Evaluation of the extract’s effect on bacterial H+-ATPase proton pumps

The ethanolic extract’s ability to inhibit H+-ATPase proton pumps in *S. aureus* ATCC 6835 was evaluated by monitoring medium acidification, following Guefack Fofack et al. [23] with modifications. Bacterial cultures underwent glucose starvation, pH adjustment, and incubation with extract at MIC and 2x MIC. Glucose addition initiated medium acidification, and pH was measured every 10 minutes for 60 minutes.

### Preparation of cream based on ethanolic extract of *Uvaria chamae*

Following Zinsou et al. [24] with modifications, the cream was prepared under hygienic conditions to ensure microbiological quality. The active ingredient was the ethanolic extract of *Uvaria chamae* roots, with sunflower oil as the main excipient, using phase inversion emulsification. Various tests were performed to assess cream quality.

### Evaluation of acute irritant and corrosive effect on skin

Dermal toxicity of the cream was assessed according to OECD guideline 404 [25]. After applying 0.5 g of cream to the skin of test rats for four hours, signs of erythema and edema were observed and scored at intervals (1, 24, 48, 72 hours) post-removal, using the Draize scale.

### *In vivo* evaluation of antibacterial effect and wound-healing potential

Wistar albino rats, acclimatized for two weeks, were divided into five batches (n=3). *S. aureus*-infected lesions were induced and treated with *Uvaria chamae* cream, following Dougnon et al. [26].

#### Lesion induction

Rats were anesthetized with ketamine hydrochloride (80 mg/kg), and a 2.25 cm² wound was excised on the left flank.

#### Lesion infection

A bacterial suspension of 10⁶ CFU/ml was applied to the lesions. Infected wounds were confirmed by swabbing and culturing on Chapman agar.

#### Treatment with *Uvaria chamae* cream

Treatment started 48 hours post-infection and continued daily for 21 days. Batches were treated as follows:

> Batch 1: Negative control (no infection, no treatment)

> Batch 2: Infected, untreated

> Batch 3: Infected, treated with 0.5 g fucidin 2% cream

> Batch 4: Uninfected, treated with 0.5 g experimental cream

> Batch 5: Infected, treated with 0.5 g experimental cream

#### Evaluation of parameters

Wounds were swabbed and cultured on Chapman agar every four days to monitor bacterial presence. Lesion progression was recorded photographically and by tracing on transparent paper. Wound retraction percentage was calculated using the formula by Kongstad et al. [27]. Wound retraction (%) = (Initial diameter-healed wound diameter)/(Initial diameter)×100

### Statistical analysis

The results of the antibacterial and wound-healing tests were analyzed using STATA 11 software. Quantitative variables were presented as mean ± standard deviation. Additionally, the means and standard deviations of the inhibition diameters for the extract were compared with those of the reference discs, using a significance level of 5%. Curves were generated using Microsoft Excel.

## Results

### Antibacterial properties of ethanolic extracts of *Uvaria chamae*

The study of the antibacterial activity of ethanolic extracts of *Uvaria chamae* demonstrated effectiveness solely against Gram-positive cocci strains. Activity was observed in all Gram-positive cocci strains tested, with average inhibition diameters ranging from 24 mm to 30 mm. The extract exhibited the highest activity against clinical strains of *Enterococcus faecalis* and methicillin-resistant *S. aureus*. The extract showed a significantly higher inhibition diameter than vancomycin for *S. aureus* strains ATCC 6835; *S. aureus* ATCC 25923; 43/SARM; 67/SARM; 04/ E. faecalis (p < 0.05). Similarly, the inhibitory effect of this extract was significantly better than that of ciprofloxacin for the 04/ E. faecalis strain (p < 0.05) (Fig 1). The minimum inhibitory concentration (MIC) was con-sistent across all bacterial strains tested at 0.125 mg/ml, while the minimum bactericidal concentrations (MBC) ranged from 0.125 to 0.25 mg/ml. With a potency index (PA) varying between 1 and 2, the ethanolic extract of *Uvaria chamae* roots exhibited bactericidal activity against the tested strains (Table 1).

**Fig 1.**
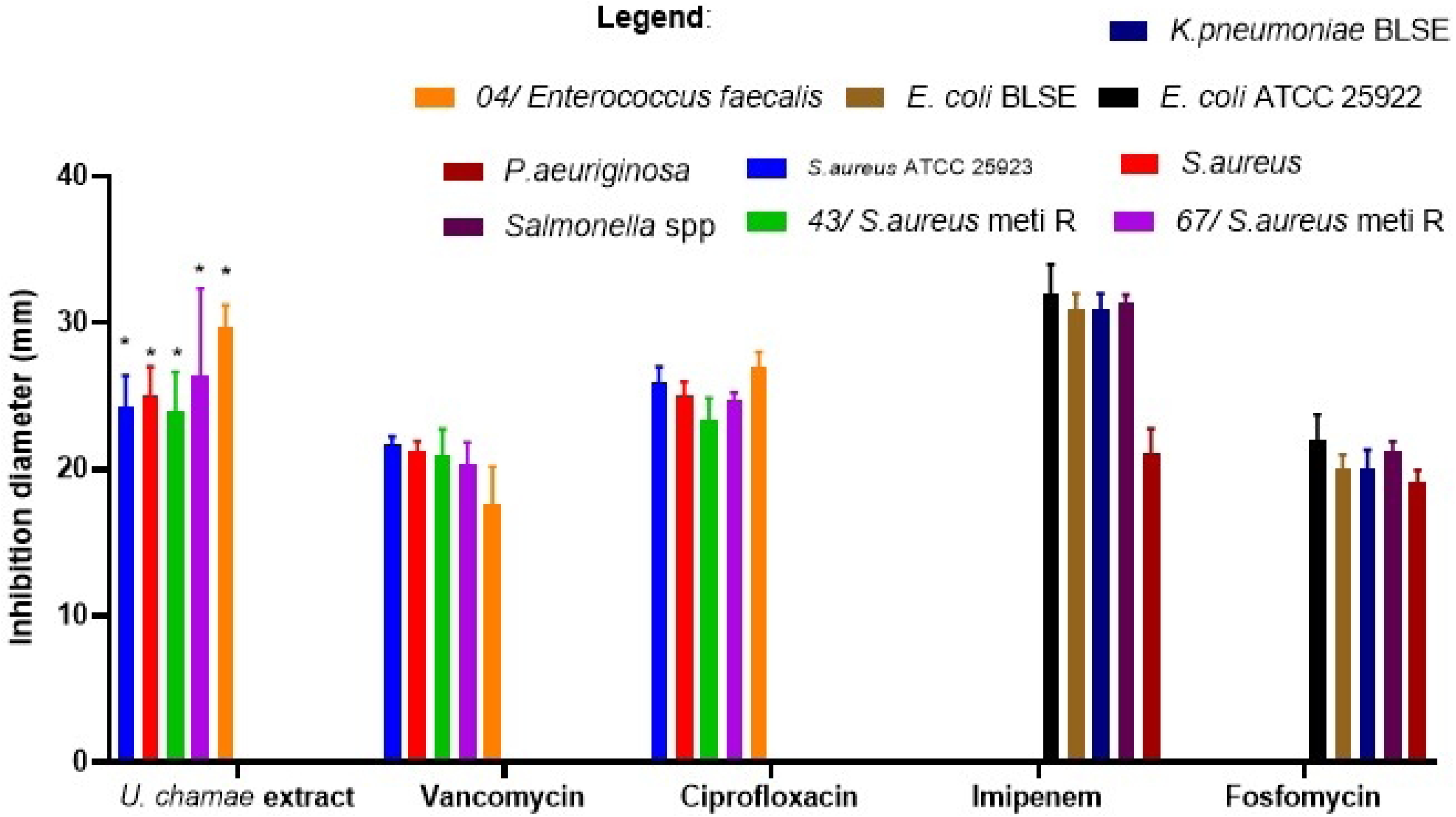
Inhibition diameter of the ethanolic extract of *Uvaria chamae* and reference antibiotics on bacterial strains.

**Table 1.**
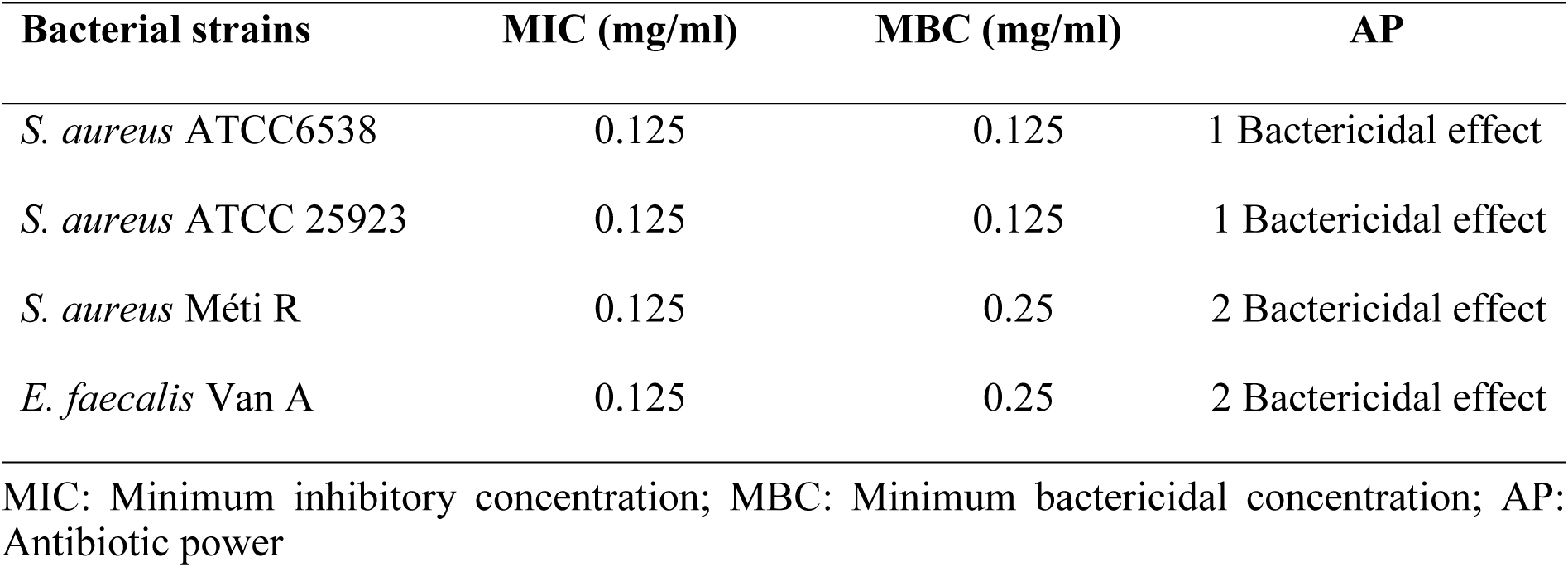
MIC, MBC and AP of the ethanolic extract of the roots of *Uvaria chamae* on the bacterial strains.

### Exploration of the mode of action of ethanolic extracts of *Uvaria chamae* on the bacterial membrane

The results of the Time Kill Kinetic Test indicated that, in the absence of treatment (negative control), the bacteria experienced exponential growth over 24 hours. However, the presence of the ethanolic extract resulted in a reduction in bacterial growth, proportional to the concentration of the extract. At various concentrations, the lag phase lasted 2 hours, followed by a stable bacterial count until the 8th hour (for MIC and 2MIC) and the 6th hour (for 4MIC). After 24 hours, the bacterial count in the medium became very low, comparable to the positive control. These findings confirm the bactericidal effect of the ethanolic extract of *Uvaria chamae* root (Fig 2).

**Fig 2.**
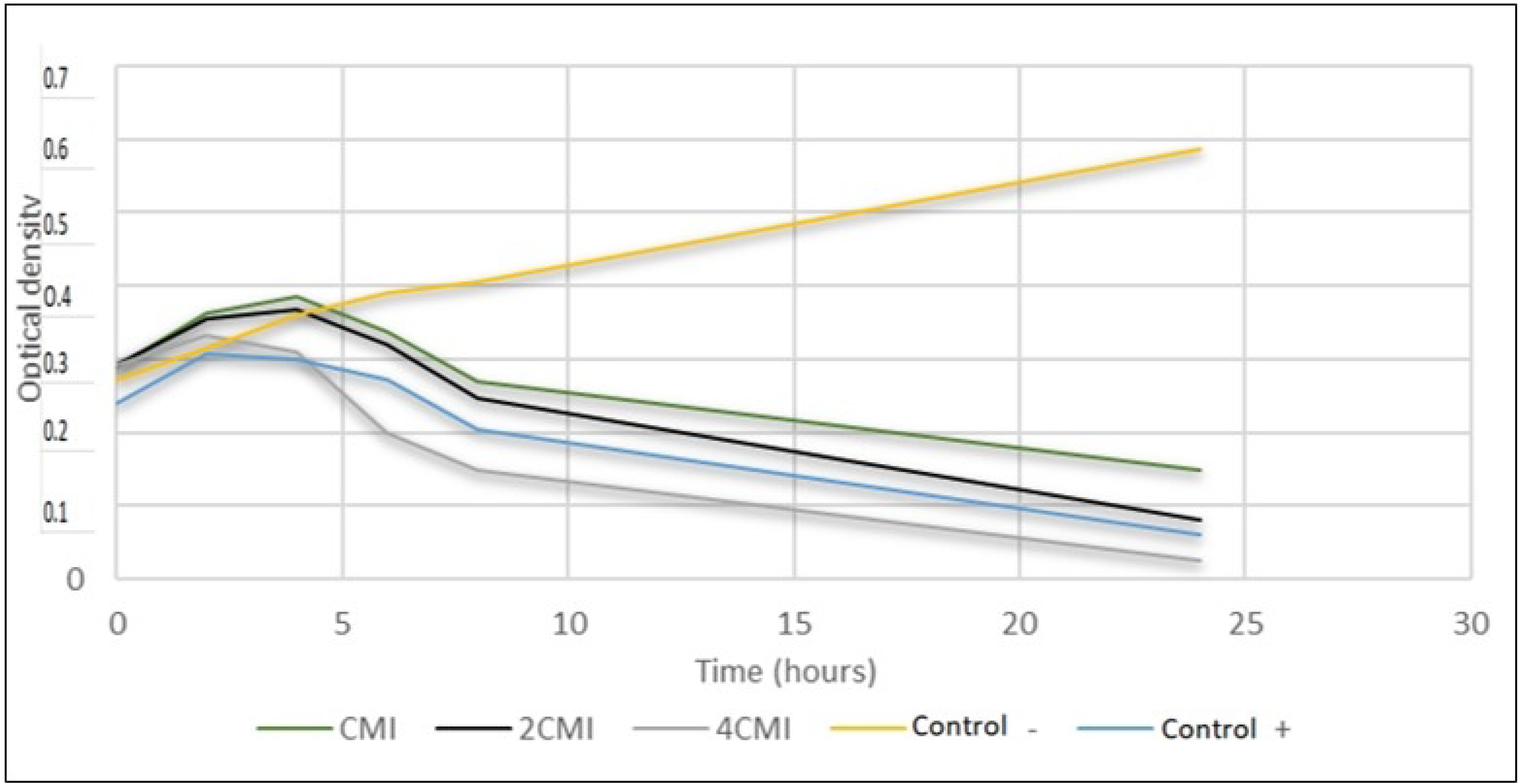
Effect of the ethanolic extract of the root of *Uvaria chamae* on the growth curve of *S. aureus*.

The study on the permeability of the outer membrane to erythromycin revealed that the ethanolic extract of *Uvaria chamae* root affected bacterial growth in the presence of erythromycin. At reduced concentrations, erythromycin exhibited a partial inhibitory effect on bacterial growth. When combined with the extract, the inhibitory activity of erythromycin on the outer membrane of *S. aureus* was enhanced, increasing bacterial sensitivity to erythromycin. These results suggest that the *Uvaria chamae* root extract disrupts outer membrane permeability (Fig 3).

**Fig 3.**
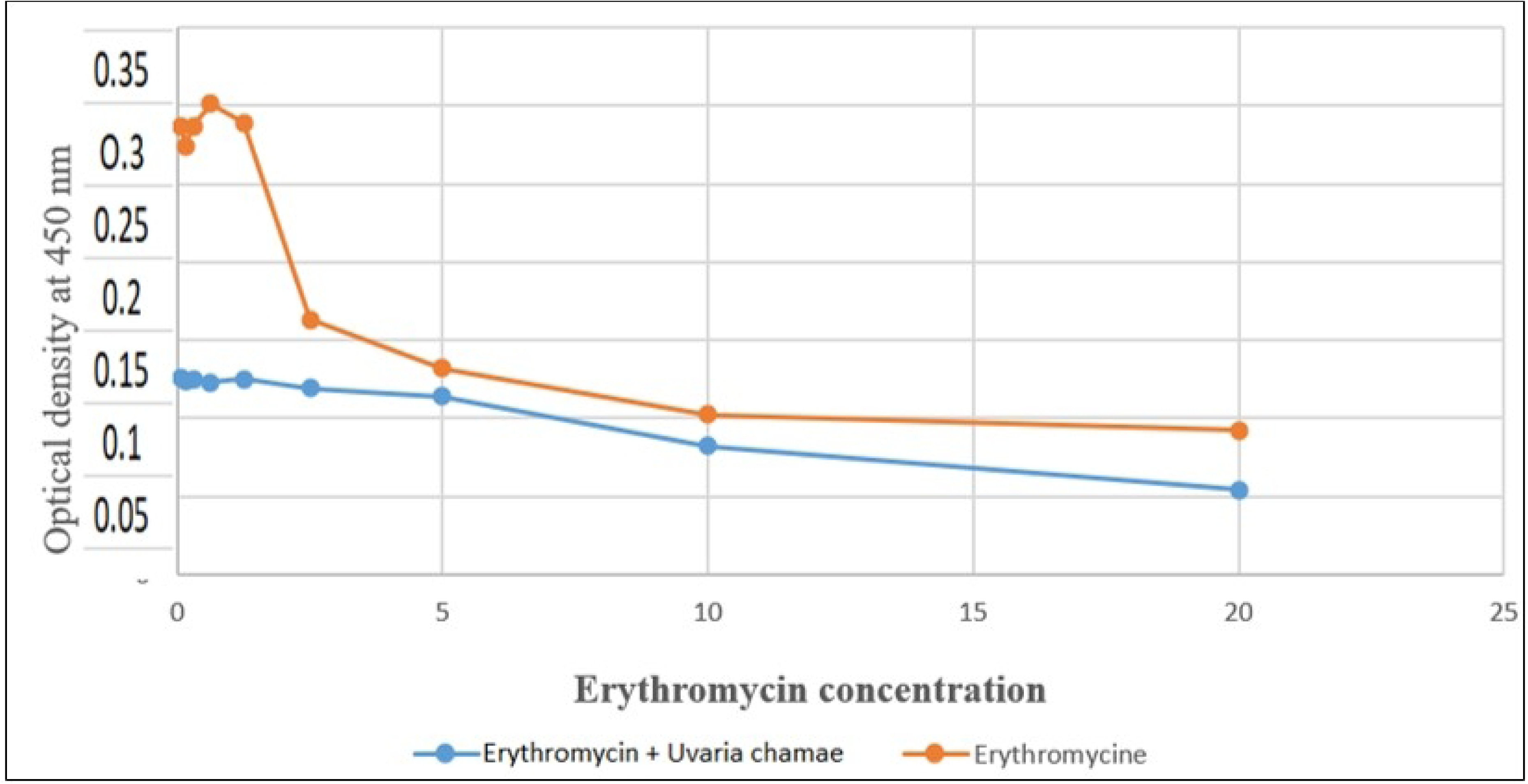
Effect of the ethanolic extract of the root of *Uvaria chamae* in combination with erythromycin on the permeability of the outer membrane of *S. aureus*.

The effect of the ethanolic extract of *Uvaria chamae* on the bacterial H+-ATPase proton pumps of *S. aureus* ATCC 6538 showed that, during the experiment, pH values changed from 6.32 to 5.72 in the presence of 3% DMSO. However, when exposed to the extract at 2MIC, pH values slightly increased from 6.32 to 6.44, while at MIC, they remained relatively stable, varying from 6.32 to 6.37. Lowering pH values promotes bacterial growth, whereas increasing pH values inhibits the growth of the studied bacteria (Fig 4).

**Fig 4.**
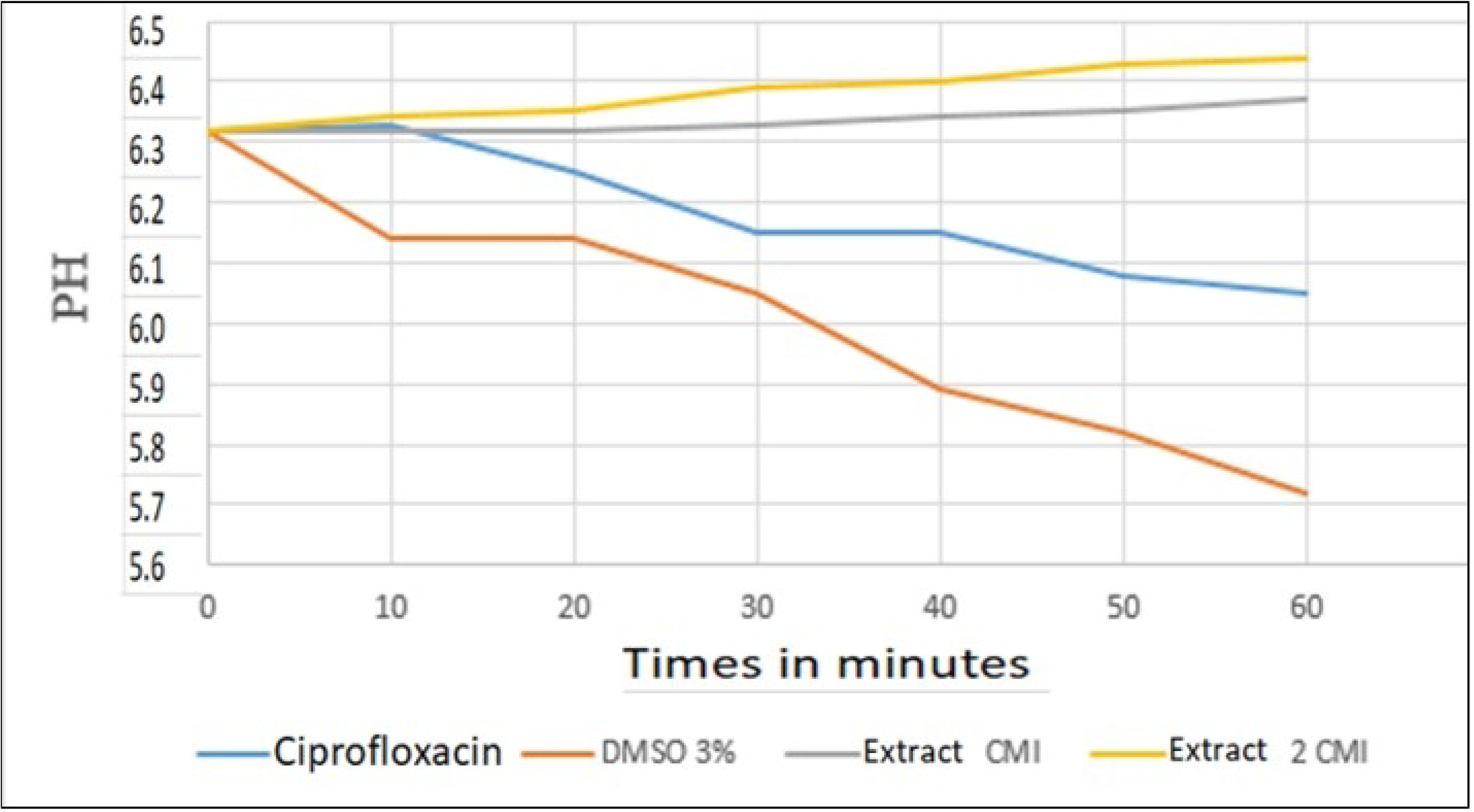
Impact of ethanolic extract of *Uvaria chamae* roots on bacterial H+-ATPase mediated by proton pumps of *S. aureus* ATCC 6538.

### Macroscopic and physicochemical characteristics of the cream based on ethanolic extract of *Uvaria chamae*

The cream formulated with ethanolic extract of *Uvaria chamae* was evaluated for its macroscopic and physicochemical properties. The results indicated that the cream had an acidic pH, good homogeneity, and a thick, smooth, shiny appearance with a soft and silky texture, making it easy to spread (Table 2). Microscopic evaluation 24 hours after formulation showed a homogeneous distribution of small oil globules, confirming the cream’s homogeneity and stability. The centrifugation test at different temperatures (4°C, 25°C, and 40°C) showed no separation after centrifugation, and the cream remained stable over 7 days of observation. Macroscopic observation of the cream kept at room temperature in the laboratory for several weeks showed no signs of instability (Table 3). Sensoro-rheological tests confirmed these findings, with participants noting a characteristic odor, soft color, fine texture, and ease of spreading (Table 4).

**Table 2.** Physico-chemical characteristics of the cream.

| Settings | Features |
| --- | --- |
| Color | Brown |
| Smell | Characteristic |
| Spreading | Easy |
| Texture | Creamy, thick, smooth appearance, shiny with a soft and silky touch |
| Consistency | Soft |
| Homogeneity | Good |
| pH | 5 |
| Viscosity | 29,035.41 mPa.s |

**Table 3.**
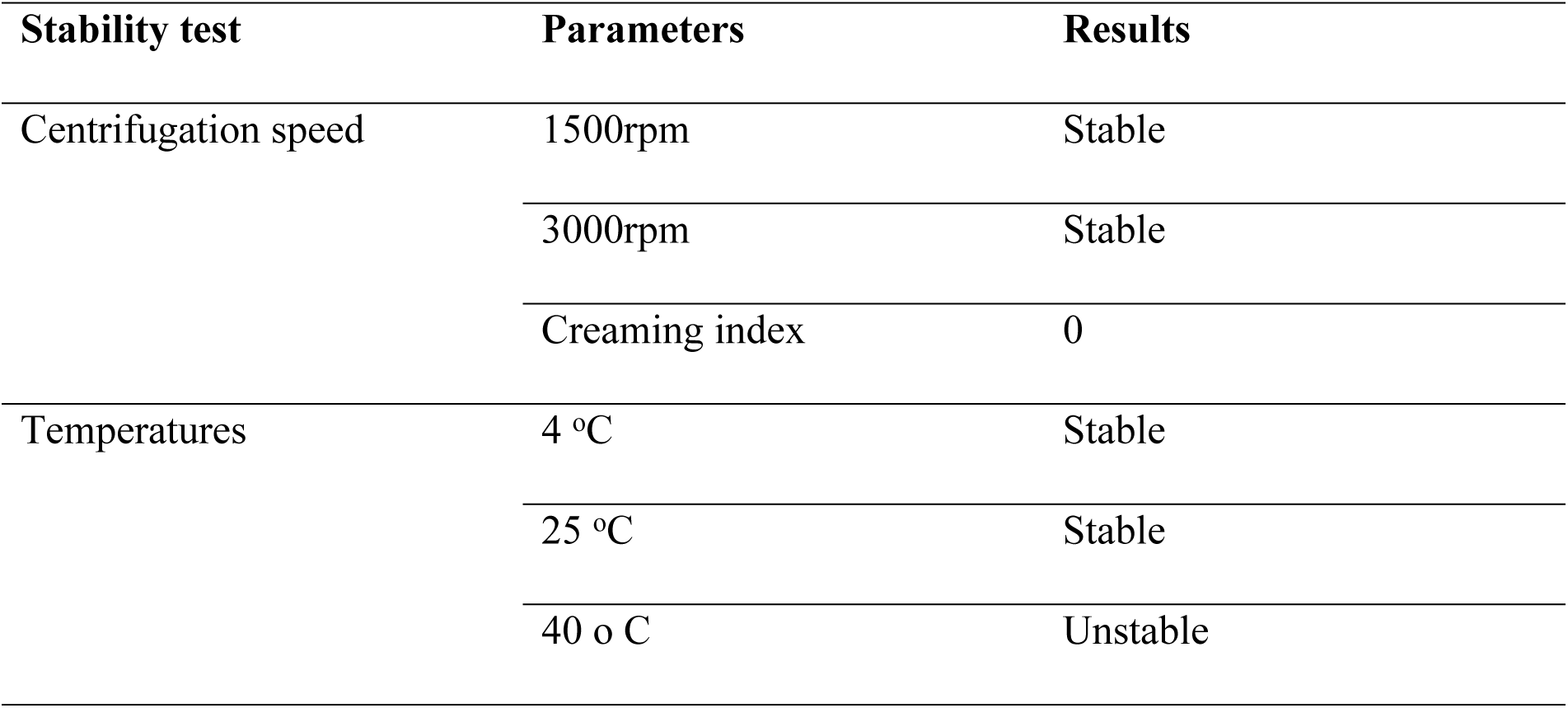
Study of the stability of the cream.

| Stability test | Parameters | Results |
| --- | --- | --- |
| Centrifugation speed | 1500rpm | Stable |
|  | 3000rpm | Stable |
|  | Creaming index | 0 |
| Temperatures | 4 °C | Stable |
|  | 25 °C | Stable |
|  | 40 o C | Unstable |

**Table 4.** Results of the sensoro-rhelogical test.

| Features | Appreciation | Response frequency |
| --- | --- | --- |
| Smell | Other | - |
|  | Features | 100% |
| Color | Other | - |
|  | Brown | 100% |
| Consistency | Compact | 12.5% |
|  | Soft | 87.5% |
| Texture | Grainy | - |
|  | Fine | 100% |
| Spreading | Difficult | - |
|  | Easy | 100% |
| Sticky effect | + | 87.5% |
|  | - | 12.5% |
| Skin irritation | + | 100% |
|  | - | - |

### Antibacterial activity against *S. aureus* and skin toxicity of cream formulated with *Uvaria chamae*

The cream’s antibacterial activity was tested against *S. aureus* ATCC 3835. At a 10% concentration, the cream produced an inhibition diameter of 8 mm. Skin toxicity tests indicated that the cream was non-irritating; none of the test rats showed signs of erythema or edema, and all survived the 14-day observation period without any significant weight differences compared to the control group (Fig 5). No significant difference was observed between the weight of the rats before application of the cream and 14 days afterwards between the rats in the control group and the rats in the test group. (p>0.05) (Table 5). The *in vivo* antibacterial activity study showed that after 9 days of treatment, no bacterial growth was observed in wounds treated with the experimental cream, while the positive control group treated with commercial Fuscidin cream showed similar results after 13 days (fig 6). The negative control group remained positive for bacterial presence until healing. The treated group’s wound diameter significantly decreased over the days, achieving complete healing by the thirteenth day with a shrinkage percentage estimated at 99.99% (Fig 7; Table 6).

**Fig 5.**
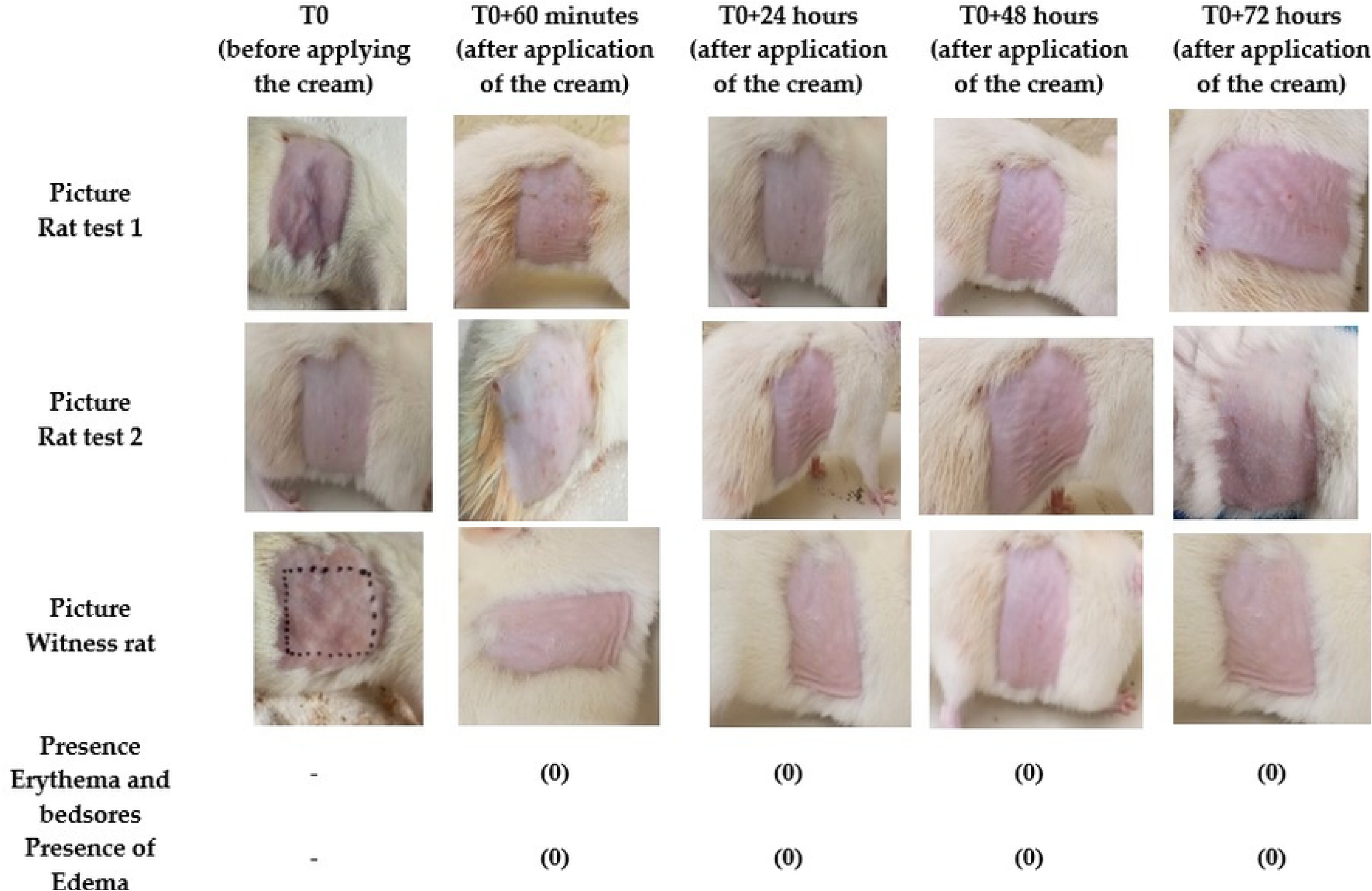
Photographic images of control and test rats in the skin toxicity test.

**Fig 6.**
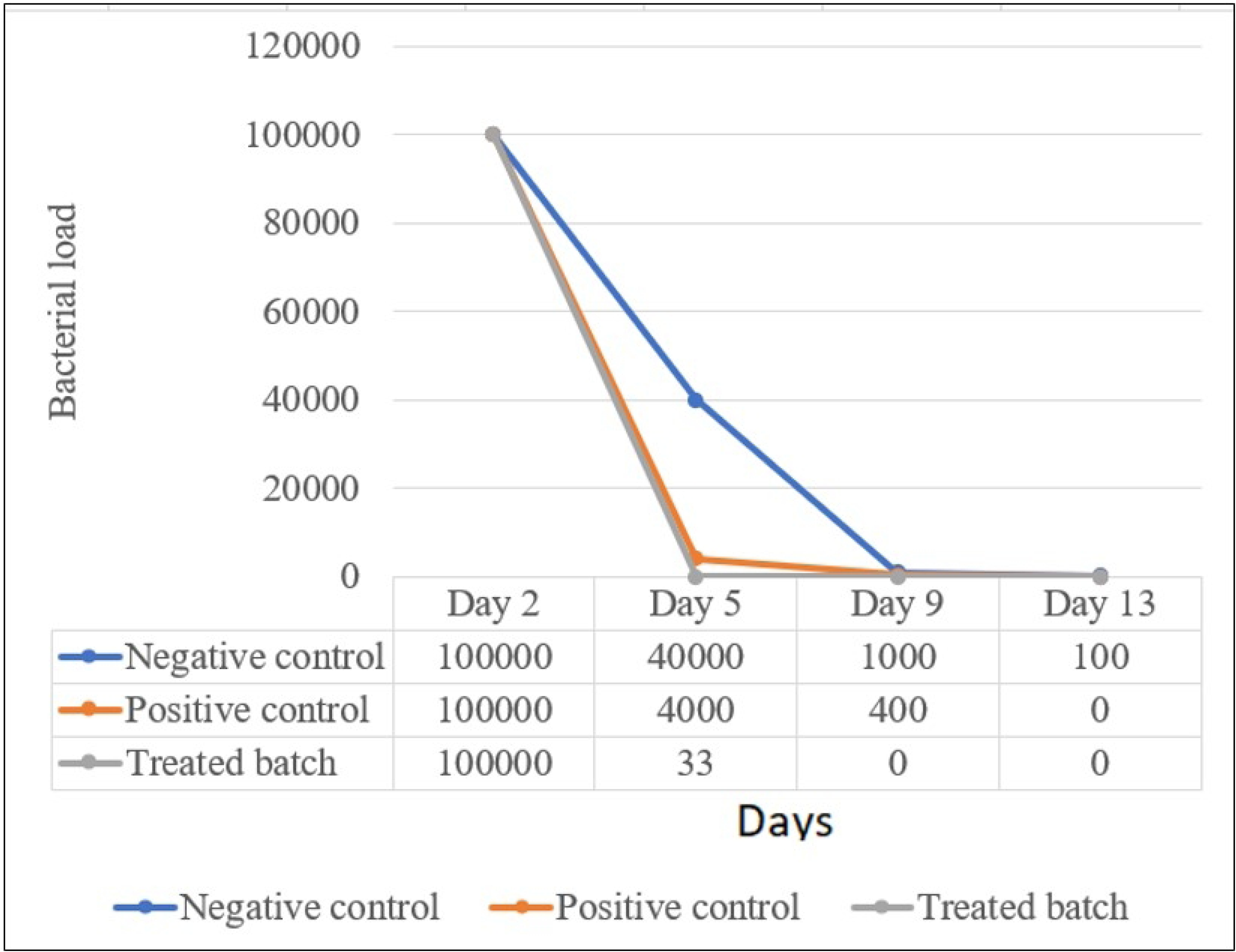
Evolution of the bacterial load of *S. aureus* ATCC 6835 as a function of time.

**Fig 7.**
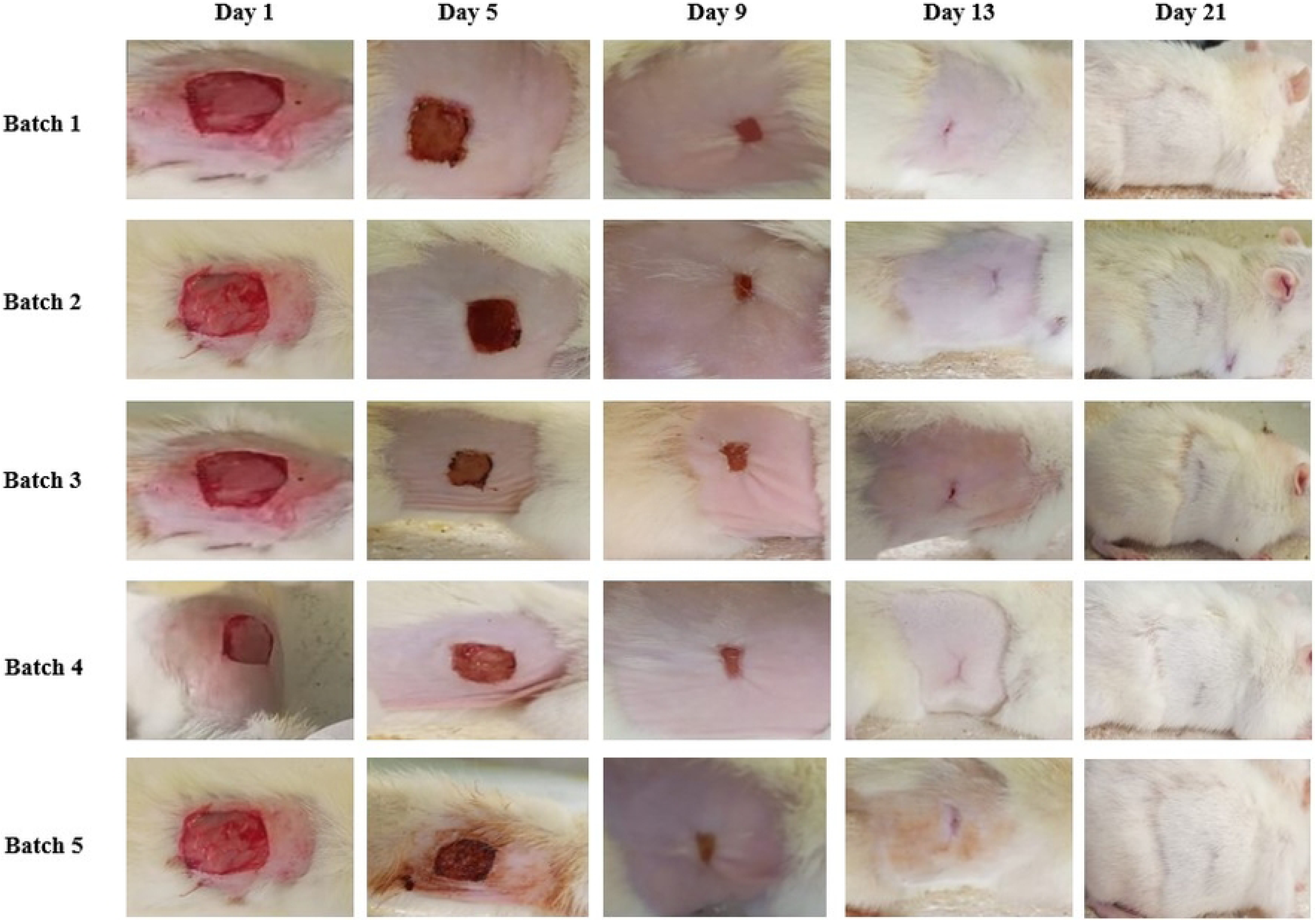
Photographic images showing impact of the cream on the surfaces of excised wounds.

**Table 5.** Evolution of body weight of rats.

|  | Control rat | Rat test | P value |
| --- | --- | --- | --- |
| Shaving weight (g) | 174.5 ± 60.10 | 189 ± 33.05 | 0.7586 |
| Weight on day 14 ( g) | 188 ± 48.08 | 200.33 ± 32.01 |  |

**Table 6.** Impact of the cream on the refraction rate (%) of excised wounds.

|  | Day 1<br>% | Day 5<br>% | Day 9<br>% | Day 13<br>% |
| --- | --- | --- | --- | --- |
| Negative control | 0 | 58.63 ± 3.61 | 91.85 ± 1.05 | 100 |
| Batch processed | 0 | 58 ± 11.26 | 94.97 ± 0.43 | 100 |
| P value | - | 0.8810 | 0.0174 | - |

## Discussion

Skin infections remain one of the most prevalent public health problems worldwide, presenting constant challenges in terms of treatment [1]. In this context, continued research to identify effective and safe antimicrobial agents is crucial for improving therapeutic strategies. The present work aims to propose a dermal formulation based on *Uvaria chamae* for the treatment of *S. aureus* skin infections.

Evaluation of the antimicrobial activity of the ethanolic extract of *Uvaria chamae* roots showed antibacterial activity according to the standards described by Tsirinirindravo and Andrianarisoa [19]. Activity was observed exclusively on the Gram-positive cocci strains tested, with an average inhibition diameter ranging between 24 mm and 30 mm. These results are consistent with the findings of Koudokpon et al. [11], who demonstrated that the selectivity of the extract towards Gram-positive cocci is due to the presence of chalcones and dihydrochalcones, which exhibit significantly higher activity on Gram-positive cocci compared to Gram-negative bacilli [28]. The bactericidal activity of the extract was demonstrated by the inhibition of bacterial growth of *S. aureus* ATCC 6538 in the presence of the extract. The antibacterial effect of the extract can be explained by the disruption of the permeability of the outer membrane of Gram-positive cocci, which are generally more sensitive because of the structure of their outer layer, rich in peptidoglycans [30]. Indeed, the outer membrane permeability test with erythromycin showed that erythromycin, at a reduced concentration, partially inhibits bacterial growth. However, the presence of the extract enhanced the permeability of the outer membrane of *S. aureus*, thereby increasing their sensitivity to erythromycin. These results are consistent with previous studies on the methanolic extract of *Persea americana* [31], which showed that the antibacterial activity of the extract on *S. aureus* was due to the disruption of the permeability of the outer membrane.

The effect of the extract on bacterial H+-ATPase proton pumps is crucial for understanding the underlying mechanisms of its antibacterial activity. H+-ATPase proton pumps are essential components of the bacterial membrane, playing a central role in maintaining intracellular pH and regulating membrane potential [30]. When impaired or inhibited, these pumps can disrupt ionic homeostasis, leading to devastating consequences for bacterial survival and function [32]. In the present study, the extract, at a concentration of 2MIC, caused a slight increase in pH from 6.32 to 6.44. This effect is concentration-dependent, and at higher concentrations, the effect would be more pronounced. These results can be explained by the fact that certain constituents of the extract could interact with the bacterial membrane, modifying its structure and indirectly affecting the integrity and function of the proton pumps [27]. Similar results were reported by Ekom and Kuete [22], who showed that the bacteriostatic and bactericidal activities of the methanol extract of *Capsicum annuum* fruits are due to the inhibition of ATPase/H+ proton pumps.

Evaluating the physicochemical characteristics of a cream is of paramount importance in the development and use of dermal products. These characteristics provide crucial information about the stability, effectiveness, and safety of the product. Texture, viscosity, dispersion, and thermal stability are physicochemical parameters that directly influence the quality and user experience of the cream [33]. The test results to assess the pharmaceutical quality of the cream are encouraging, showing excellent macroscopic parameters and satisfactory homogeneity. The pH of the cream is 5, which complies with cosmetic standards suitable for skin application [34]. The uniform distribution of oil globules and the stability test confirm the stability of the cream at 4°C and 25°C, in agreement with the results reported by Banerjee et al [35]. Sensory evaluations carried out with volunteers confirmed the marketability of the cream, highlighting characteristics such as pleasant consistency, characteristic color, pleasant odor, and a homogeneous, creamy, fine, and smooth appearance.

Evaluating the antibacterial activity of a cream is a critical step to ensure its therapeutic effectiveness in treating skin infections. The antibacterial effectiveness of the formulated cream was studied at different concentrations, showing a dose-dependent effect with an inhibition diameter of 8 mm at 10%. However, vancomycin incorporated into the base cream exhibited significant antibacterial activity at lower concentrations, revealing dose-dependent efficacy like that reported by Bene et al. [36]. Tolerance to a product refers to the ability of an individual, system, or organism to accept or tolerate that product without experiencing harmful or undesirable effects [31]. The skin tolerance of the cream was evaluated, and no irritant reaction was observed according to the Draize scale. This can be attributed to the non-irritating nature of the base cream and previous studies demonstrating the non-cytotoxicity of *Uvaria chamae* extracts to human cells [37–40].

Healing, defined as the shrinkage of an injured area [29], was experimentally confirmed by the cream containing the ethanolic extract of the root of *Uvaria chamae,* exhibiting remarkable healing activity. The cream induced a significant increase in the refraction rate of the wound (94.97% on the 9th day) and resulted in complete elimination of the infection (CFU of *S. aureus*) after five days of treatment. These results suggest that the ethanolic extract of *Uvaria chamae* root improves wound healing in addition to its antibacterial properties. This healing action is attributed to the phytochemicals present in the extract, such as flavonoids, polyphenols, and tannins, as reported in previous studies [11,14,41]. These results validate the use of *Uvaria chamae* extracts in the management of *S. aureus* skin infections without adverse consequences. These findings pave the way for the preclinical trial phase with a view to placing this cream on the market, offering promising prospects for its potential effectiveness in the treatment of skin infections.

This study demonstrated the antibacterial and wound-healing efficacy of the ethanolic extract of *Uvaria chamae* roots. The extract showed bactericidal activity against Gram-positive cocci, enhanced the permeability of erythromycin, and inhibited bacterial ATPase/H+ proton pumps. The cream formulation displayed excellent physicochemical properties and stability, confirmed by sensory evaluations. It showed significant antibacterial activity and remarkable wound-healing properties, making it a promising candidate for treating skin infections. Further clinical studies are recommended to confirm its efficacy in real-life applications.

## Conclusion

This study highlights the potential of *Uvaria chamae* as a valuable resource in the management of *S. aureus* skin infections. The ethanolic extract of *Uvaria chamae* roots has shown significant antibacterial properties against a range of *S. aureus* strains, demonstrating bactericidal effects by permeability of the bacterial membrane. The formulation of a cream based on this extract proved stable, with favorable physicochemical characteristics and excellent sensory appeal. The cream showed significant antibacterial activity against S. aureus, while demonstrating non-irritant properties in skin toxicity tests. In addition, the cream accelerated wound healing, demonstrating its dual efficacy in fighting infection and promoting tissue repair. These results not only contribute to the growing body of evidence supporting *Uvaria chamae* antimicrobial potential, but also underline its suitability for topical application in the treatment of bacterial skin infections.

## Acknowledgments

The authors are grateful to all the staff of the Research Unit in Applied Microbiology and Pharmacology of natural substances involved in this study.

## Supporting information

**S1_Fig 1. Inhibition diameter of the ethanolic extract of *Uvaria chamae* and reference antibiotics on bacterial strains**

**S2_Fig 2. Effect of the ethanolic extract of the root of *Uvaria chamae* on the growth curve of *S. aureus***

**S3_Fig 3. Effect of the ethanolic extract of the root of *Uvaria chamae* in combination with erythromycin on the permeability of the outer membrane of *S. aureus***

**S4_Fig 4. Impact of ethanolic extract of *Uvaria chamae* roots on bacterial H+-ATPase mediated by proton pumps of *S. aureus* ATCC 6538**

**S5 _Fig 5. Photographic images of control and test rats in the skin toxicity test.**

**S6_Fig 6. Evolution of the bacterial load of *S. aureus* ATCC 6835 as a function of time.**

**S7_Fig 7. Photographic images showing impact of the cream on the surfaces of excised wounds.**

## References

1. Yakupu A, Aimaier R, Yuan B, Chen B, Cheng J, Zhao Y, et al. The burden of skin and subcutaneous diseases: findings from the global burden of disease study 2019. Front Public Health. 2023, 11, 1145513. DOI: 10.3389/fpubh.2023.1145513.

2. Francès P, Merikhi A, Halbout O, Stoumen M. Trois exemples d’infections bactériennes cutanées. Aide-Soignante. 2023, 37, 24–7. 10.1016/j.aidsoi.2023.02.008.

3. Burillo A, Pulido-Pérez A, Bouza E. Current challenges in acute bacterial skin infection management. Curr Opin Infect Dis. 2024, 37, 71–9. DOI: 10.1097/QCO.0000000000000989.

4. Larquey M, Mahé E. Infections cutanées à staphylocoque et streptocoque chez l’enfant. Perfect En Pédiatrie. 2018, 1, 25–31. 10.1016/j.perped.2018.01.015.

5. Socohou A, Sina H, Degbey C, Adjobimey T, Sossou E, Boya B, et al. Pathogenicity and Molecular Characterization of *Staphylococcus aureus* Strains Isolated from the Hospital Environment of CHU-Z Abomey-Calavi/Sô-Ava (Benin). BioMed Res Int. 2021, e6637617. DOI: 10.1155/2021/6637617.

6. Bernard P. Pour une meilleure prise en charge des infections bacteriennes courantes. Ann Dermatol Vénéréologie. Elsevier. 2019, 1–5. DOI: 10.1016/j.annder.2019.05.008

7. Bernabé KJ, Langendorf C, Ford N, Ronat J-B, Murphy RA. Antimicrobial resistance in West Africa: a systematic review and meta-analysis. Int J Antimicrob Agents. 2017, 50, 629–39. DOI: 10.1016/j.ijantimicag.2017.07.002

8. Cowan MM. Plant Products as Antimicrobial Agents. Clin Microbiol Rev. 1999, 12, 564–82. DOI: 10.1128/CMR.12.4.564.

9. Salam AM, Quave CL. Opportunities for plant natural products in infection control. Curr Opin Microbiol. 2018, 45:189–194. DOI: 10.1016/j.mib.2018.08.004

10. Koudokpon H, Dougnon VT, Bankolé HS, Fah L, Hounmanou YMG, Baba-Moussa L. et al. Enquête ethnobotanique sur les plantes utilisées dans le traitement des infections au Sud-Bénin. Health Sci Dis. 2017, 18, 1–8. 10.5281/hsd.v18i2.809.

11. Koudokpon H, Armstrong N, Dougnon TV, Fah L, Hounsa E, Bankolé HS, et al. Antibacterial Activity of Chalcone and Dihydrochalcone Compounds from *Uvaria chamae* Roots against Multidrug-Resistant Bacteria. BioMed Res Int. 2018, 1453173. DOI: 10.1155/2018/1453173

12. Daï EH, Houndonougbo JS, Idohou R, Ouédraogo A, Kakaï RG, Hotes S, et al. Modeling current and future distribution patterns of *Uvaria chamae* in Benin (West Africa): Challenges and opportunities for its sustainable management. Heliyon. 2023, 9, e13658, 1–11. DOI: 10.1016/j.heliyon.2023.e13658

13. Dougnon V, Legba B, Koudokpon H, Klotoe JR, Bankole H, Dougnon J. Bioactives and Pharmacology of Uvaria chamae P. Beauv. Bioact Pharmacol Med Plants. 2022. (1st ed., pp. 1–8). Apple Academic Press. DOI: 10.1201/9781003281658-5

14. Ochiabut OM-TB, Unaeze B.C, Chukwura EI, Uzoewulu NG. Phytochemistry and Antimicrobial Activities of Crude and Diluted Leaves and Roots Extracts of *Uvaria chamae* on Selected Bacteria. J Pharm Res Int. 2022, 34, 7–18. DOI: 10.9734/JPRI/2022/v34i8A35473

15. Bentabet N, Rajaa R, Sakina N. Enquête ethnobotanique et inventaire des plantes médicinales utilisées dans le traitement des maladies dermatologiques dans la ville d’Ain Temouchent. J Appl Biosci. 2022, 1, 17704– 17719. DOI: 10.35759/JABs.170.4

16. Benhiba H, Hassam B. Phytophotodermatose à la *Mentha rotundifolia*. Pan Afr Med J. 2014, 18, 192. DOI: 10.11604/pamj.2014.18.192.4923

17. Agbankpe AJ, Dougnon TV, Bankole SH, Houngbegnonn O, Dah-Nouvlessouno D, Baba-Moussa L. (). In vitro antibacterial effects of Crateva adansonii, Vernonia amygdalina and Sesamum radiatum used for the treatment of infectious diarrhoeas in Benin. J Infect Dis Ther 20164: 281–287. DOI:10.4172/2332-0877.1000281

18. CASFM. Comité de l’Antibiogramme de la Société Française de Microbiologie. Société Fr. Microbiol. 2023. https://www.sfm-microbiologie.org/wp-content/uploads/2023/06/CASFM2023_V1.0.pdf

19. Tsirinirindravo L, Andrianarisoa B. Activités antibactériennes de l’extrait des feuilles de *Dalechampia clematidifolia* (Euphorbiaceae). Int J Biol Chem Sci. 2009, 5, 1198–1202. DOI: 10.4314/ijbcs.v3i5.51098

20. Dougnon V, Hounsa E, Koudokpon H, Agbodjento E, Afaton A, Sintondji K, et al. Assessment of the antibacterial effect of *Khaya senegalensis* on some Gram-negative bacteria. Bull Natl Res Cent. 2021, 45, 107, 1–8. 10.1186/s42269-021-00568-0.

21. Djague F, Lunga PK, Toghueo KRM, Melogmo DYK, Fekam BF. *Garcinia kola* (Heckel) and *Alchornea cordifolia* (Schumach. & Thonn.) Müll. Arg. from Cameroon possess potential antisalmonella and antioxidant properties. PLoS ONE. 2020, 8, 1–15. DOI: 10.1371/journal.pone.0237076

22. Ekom SE, Tamokou J-D-D, Kuete V. Antibacterial and Therapeutic Potentials of the *Capsicum annuum* Extract against Infected Wound in a Rat Model with Its Mechanisms of Antibacterial Action. BioMed Res Int. 2021, 2021, 1–17. DOI: 10.1155/2021/4303902.

23. Guefack Fofack M-G, Tankeo S, Carine Marcelle N, Nayim P, Wamba B, Bonsou I, et al. Antibiotic-potentiation activities of three animal species extracts, *Bitis arietans*, Helix aspersa, and *Aristaeomorpha foliacea* and mode of action against MDR Gram-negative bacteria phenotypes. Investig Med Chem Pharmacol. 2021, 4, 1–15. DOI: 10.31183/imcp.2020.00048.

24. Zinsou A, Assanhou AG, Ganfon H, Sounouvou H, Kassehin UC, Lawson RF, et al. Development of new dermatological formulations for the treatment of cutaneous candidiasis. Sci Afr. 2020, 8, e00342. 10.1016/j.sciaf.2020.e00342.

25. OCDE. Essai n° 404 : Effet irritant/corrosif aigu sur la peau. 2015, 1–9. https://www.oecd.org/fr/publications/essai-n-404-effet-irritant-corrosif-aigu-sur-la-peau-9789264242685-fr.htm.

26. Dougnon V, Klotoé J, Bankole H, Yaya Nadjo S. Antibacterial and wound healing properties of *Terminalia superba* Engl. and Diels (Combretaceae) in albino wistar rats. J Bacteriol Parasitol. 2014, 5, 1–7. DOI: 10.4172/2155-9597.1000206.

27. Kongstad KT, Wubshet SG, Kjellerup L, Winther A-ML, Staerk D. Fungal plasma membrane H+-ATPase inhibitory activity of o-hydroxybenzylated flavanones and chalcones from *Uvaria chamae* P. Beauv. Fitoterapia. 2015, 105, 102–6. 10.1016/j.fitote.2015.06.013

28. Khan I, Rahman H, Abd El-Salam NM, Tawab A, Hussain A, Khan TA, et al. *Punica granatum* peel extracts: HPLC fractionation and LC MS analysis to quest compounds having activity against multidrug resistant bacteria. BMC Complement Altern Med. 2017, 247, 1–6. 10.1186/s12906-017-1766-4

29. Ekom SE, Tamokou JDD, Kuete V. Methanol extract from the seeds of *Persea americana* displays antibacterial and wound healing activities in rat model. J Ethnopharmacol. 2022, 282, 114573. DOI: 10.1016/j.jep.2021.114573

30. Kobayashi H. A proton-translocating ATPase regulates pH of the bacterial cytoplasm. J Biol Chem. 1985, 260, 72–6. https://www.jbc.org/article/S0021-9258(18)89694-6/pdf

31. Ivanova K, Ramon E, Ivanova A, Sanchez-Gomez S, Tzanov T. Bio-Based Nano-Enabled Cosmetic Formulations for the Treatment of *Cutibacterium acnes*-Associated Skin Infections. Antioxidants. 2023, 12, 432, 1–16. DOI: 10.3390/antiox12020432

32. Oliveira DM, Melo FG, Balogun SO, Flach A, de Souza ECA, de Souza GP, et al. Antibacterial mode of action of the hydroethanolic extract of *Leonotis nepetifolia* (L.) R. Br. involves bacterial membrane perturbations. J Ethnopharmacol. 2015, 172, 356–63. DOI: 10.1016/j.jep.2015.06.027

33. Ferarsa S, Zhang W, Moulai-Mostefa N, Ding L, Jaffrin MY, Grimi N. Recovery of anthocyanins and other phenolic compounds from purple eggplant peels and pulps using ultrasonic-assisted extraction. Food Bioprod Process. 2018, 109,19–28. 10.1016/j.fbp.2018.02.006

34. Ijaz N, Durrani AI, Rubab S, Bahadur S. Formulation and characterization of Aloe vera gel and tomato powder containing cream. Acta Ecol Sin. 2022, 42, 34–42. 10.1016/j.chnaes.2021.01.005

35. Banerjee K, Thiagarajan N, Thiagarajan P. Formulation and characterization of a *Helianthus annuus* - alkyl polyglucoside emulsion cream for topical applications. J Cosmet Dermatol. 2019, 18, 628–637. DOI: 10.1111/jocd.12756

36. Bene K, Camara D, Soumahoro IA, Kanga Y, Zirihi G. Formulation galénique d’une pommade antimicrobienne à base d’un extrait hydroalcoolique de *Bersama abyssinica* Fresen. Ethnopharmacologia. 2017, 60–9. http://www.ethnopharmacologia.org/wp-content/uploads/2014/05/Article-Bene-58.pdf

37. Awodiran MO, Adepiti AO, Akinwunmi KF. Assessment of the cytotoxicity and genotoxicity properties of *Uvaria chamae* P. Beauv (Annonaceae) and *Morinda lucida* Benth (Rubiaceae) in mice. Drug Chem Toxicol. 2018, 41, 232–237. DOI: 10.1080/01480545.2017.1365884.

38. Olumese F, Onoagbe I, Eze G, Omoruyi F. Subchronic toxicity study of ethanolic extract of *Uvaria chamae* root in rats. Trop J Pharm Res. 2018, 17, 831–836. DOI: 10.4314/tjpr.v17i5.12.

39. Legba B, Dougnon V, Deguenon E, Agbankpe J, Senou M, Aniambossou A, et al. Toxicological characterization of six plants of the beninese pharmacopoeia used in the treatment of salmonellosis. J Toxicol. 2019, 2019, 1–12. DOI: 10.1155/2019/3530659.

40. Olumese FE, Omoruyi FO, Onoagbe IO. Effects of *Uvaria chamae* root extracts on blood glucose, inflammatory markers, lipid profile, liver and renal status in streptozotocin-induced diabetic rats. Niger J Physiol Sci. 2019, 34, 207–213. https://pubmed.ncbi.nlm.nih.gov/32343272/#:~:text=Overall%2C%20the%20consumption%20of%20Uvaria%20chamae%20extracts%20lowered,effects%20on%20the%20kidney%20in%20this%20short-term%20study.

41. Enin GN, Shaibu SE, Ujah GA, Ibu RO, Inangha PG. Phytochemical and Nutritive Composition of *Uvaria chamae* P. Beauv. Leaves, Stem Bark and Root Bark. Chem Search J. 2021, 12, 9–14. https://www.ajol.info/index.php/csj/article/view/209958

